# Mandrel Diameter is a Dominating Parameter for Fiber Alignment Control in Rotating Mandrel Electrospinning Systems

**DOI:** 10.1101/2024.07.11.603153

**Authors:** Katherine L. Meinhold, Tyler Tankersley, Rylie Darlington, Jennifer L. Robinson

## Abstract

Aligned nano and micron-sized electrospun scaffolds are advantageous for 3D *in vitro* models of fibrous, aligned tissue. A common approach to induce alignment is to collect on a rotating mandrel at high rotational speeds. Historically, rotating mandrel speed has been considered the major driver in tuning the degree of alignment even though mandrel diameter is known to modulate linear velocity and increase alignment. However, the comparative impact of mandrel diameter vs. rotating mandrel speed has not been systemically investigated. As such, this study aimed to investigate the role of mandrel diameter on fiber alignment, fiber fraction, and fiber diameter under controlled modulation of common processing parameters including applied voltage, distance to collector, and mandrel rotational speed. Analysis of all samples was performed using scanning electron microscopy (SEM) and image analysis by the DiameterJ and OrientationJ plugins in ImageJ. Using linear regression analysis in JMP software, mandrel diameter was shown to be the dominant factor influencing fiber diameter, fiber fraction, and fiber alignment of samples at all tested conditions including increased rotational speed. Overall, these findings suggest that rather than increasing rotational speed of the collector, fiber alignment can be more finely tuned by increasing mandrel diameter.

**Graphical Abstract:** 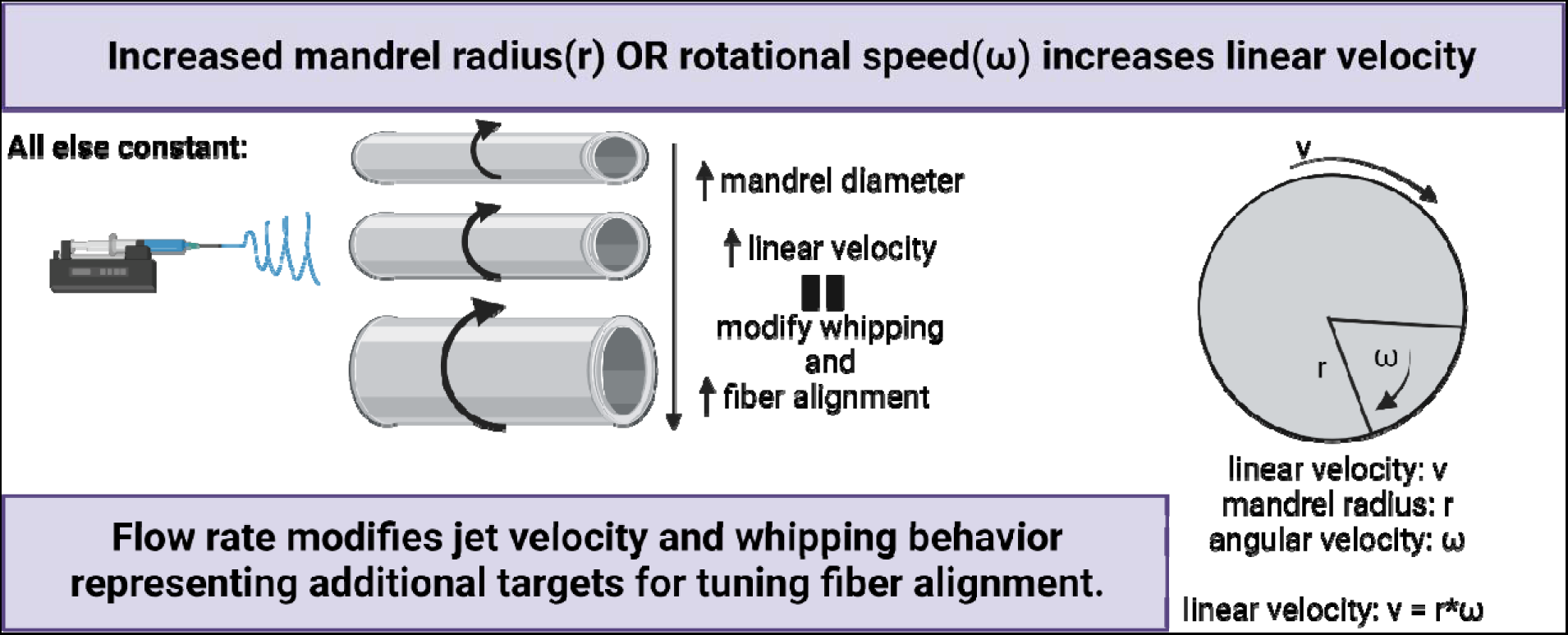

## Introduction

Fibrous materials are advantageous for use as 3D *in vitro* models of fibrous, aligned tissue as they provide a large surface area-to-volume ratio, interconnected porosity, and a wide range of mechanical properties.^1–3^ When considering applications in tissue engineering, alignment must also be considered as many native tissues with poor healing potential including connective tissues have regions of high alignment and anisotrophy.^4^ Many fabrication methods exist to obtain a nanofibrous mesh, including phase separation^1,2,5^, template synthesis^1,2^, physical drawing^1,2^, self-assembly^1,2,6^, melt blowing^2,7^, and electrospinning^1,2,8^. Electrospun scaffolds in particular have been shown to modulate cell behavior and provide superior metabolic and extracellular matrix forming potential.^4^ A basic electrospinning system consists of three parts: a high voltage power supply, a polymer extrusion method, and a collector. Compared to other techniques, electrospinning offers a high degree of control and relatively easy method for creating complex, fibrous scaffolds with mechanical and structural properties mimicking native ECM in complex fibrous tissues.^9^ The primary challenges associated with electrospinning are the extensive solution, processing, and environmental parameters that affect the process, many of which are not well controlled in traditional systems.^2,10^ Systems can be further complicated by the addition of more processing parameters in order to synthesize aligned electrospun fibers as most often modifications are made in the form of additional electrostatic steering of the fibers at the collector, mechanical methods including a rotating collector, or a combination.^11–13^

Mechanical methods to modulate alignment include rotating collectors^1,12–22^, post-drawing of fibers, and centrifugal electrospinning^23^ and electrostatic methods include gap electrospinning^12,13,20,21,23–25^, magnetic field application^12,21^, and near field electrospinning^1,13^. Each of these methods have been used successfully by researchers to obtain aligned fibers; however, each has a set of specific advantages and disadvantages which are briefly summarized for gap electrospinning, magnetic field application, near field electrospinning, and rotating collectors. Gap electrospinning leverages the use of two parallel electrodes with a void gap between them to achieve fiber alignment.^23–25^ This method creates very highly aligned nano- to micron-sized fibers and is also fairly easy to add to any lab-built electrospinning set-up. However, efficacy depends on the size of the gap, which must be relatively small, thus limiting overall surface area and sample thickness.^1^ Magnetic field electrospinning induces fiber alignment by application of a magnetic field at the collector via permanent magnets. Success is dependent on the magnetic susceptibility of the polymer, the electrospinning solution, and the strength of the magnetic field.^1^ Near-field electrospinning is a 3D printer-like technique where the whipping motion of the jet is removed by manipulation of processing parameters. This technique is very effective at creating aligned scaffolds and uses lower applied voltage compared to typical electrospinning. However, there are strict limits on flow rates and the fiber diameters are limited to a 3-6μm range.^1,26^ Rotating mandrels are attractive due to the ability to collect a sample with a relatively large surface area and ease of addition to systems.^11,24^ In this technique the collector works to induce alignment by application of a drawing force on the electrospun jet and to improve alignment of collected samples the rotating collector geometry has been modified successfully^16,23,24^, however, this generally imposes new limitations.

This study utilized a rotating mandrel due to the ease of addition and the ability to create samples with large surface area making this approach the preferred method in commercial electrospinning set-ups. The literature generally concludes that an increased rotational speed when collecting aligned fibers on a rotating mandrel leads to increased fiber alignment.^12,27–33^ This is due to the linear relationship between increased drum or mandrel diameter and linear speed and the effect of a drawing force on the bending instability of electrospun jets.^5,12^ It is considered that at a minimum the collector surface speed must match or exceed the rate of fiber production (solution flow rate) and increased linear velocity can also provide a drawing force to contribute to fiber alignment. However, there is high variability in the degree of highly aligned fibers even with very high collection speeds and this is the main parameter that is optimized to modulate fiber alignment.^11,34^ Beyond modification of collector type or the speed and geometry of rotating collectors, other studies have suggested that the key to consistently obtaining highly aligned fibers is to eliminate the characteristic bending instability.^11^ Some studies have included mandrel diameter as a factor in system optimization, however, these studies have concluded that diameter is either of limited importance^11,27^ or have not systematically explored the impact and extrapolated the meaning for wider application in collecting aligned electrospun fibers^28^.

Thus, the focus of this study was to provide a systematic investigation of the relationship between mandrel diameter and collected fiber alignment at multiples speeds, collection distances, and applied voltage. In this paper, we aim to demonstrate the key relationship between fiber diameter, fiber alignment, and each of the given processing parameters including applied voltage, collection distance, rotational speed, and mandrel diameter. The impact of rotating mandrel diameter, speed, excitation voltage, and distance to collector were modulated and resulting fiber diameter, fiber fraction, and fiber alignment were assessed. These relationships can be extrapolated for use in systems with different polymer and solvent systems and can provide a guide for designing and fabricating small-scale electrospinning systems to obtain aligned fibrous scaffolds.

## Materials and Methods

PCL (50,000 Mw) was purchased from CAPA lot # 120625. For all studies, 20% w/v PCL was dissolved in CHCl_3_. Samples were collected at high relative humidity (65%± 10), flow rate of 1.0 mL/hr, and 15 minutes at all mandrel sizes, collection distances, and applied voltages. Mandrel diameters of ¾”, 1”, and 2”, collection distances of 15cm and 30cm, applied voltages of 15kV and 18kV, and rotational speeds of 580±17 RPM, 780 RPM, and 1010±17 RPM were tested. After collection, samples were dried in a fume hood overnight and then under vacuum for 8 hours before characterization. At least three specimens were fabricated per sample group. **Figure 1** provides a visual overview of the main physically manipulated study parameters and **Figure S1** shows a graphical outline of all parameters examined within the study separated into groups by mandrel diameter.

**Figure 1.**
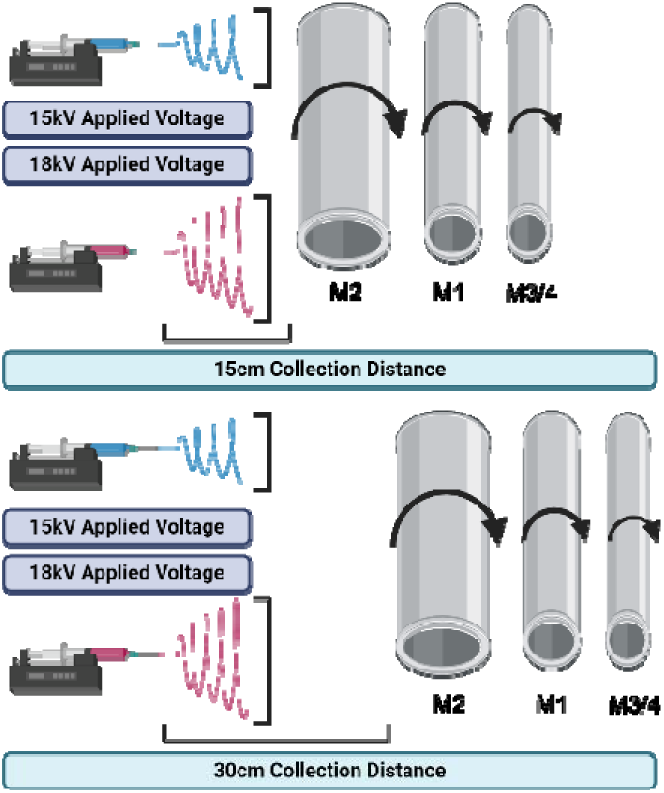
Overview of manipulated processing parameters.

Rotational speeds were determined by filming rotating mandrels with a high-speed camera at a set engine power and determining number of rotations per second at each mandrel diameter. **Table 1** shows the conversion of rotational speed to linear speed for each low, medium, and high speed used with each mandrel diameter. All conversions were performed using **Equations 1–2**., shown below.

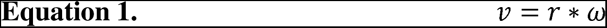

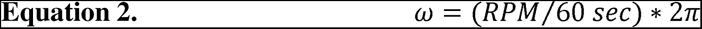

**Table 1.** Conversion of rotational speed to linear speed for ¾, 1, and 2” mandrel diameters.

| Mandrel Diameter<br>(in) | Rotational Speed<br>(RPM) | Linear Speed |  |
| --- | --- | --- | --- |
|  |  | (in/s) | (m/s) |
| ¾ | 580±17 | 46±1 | (1.16±0.03) |
| ¾ | 780 | 61 | (1.56) |
| ¾ | 1010±17 | 79±1 | (2.01±0.03) |
| 1 | 580±17 | 61±2 | (1.54±0.05) |
| 1 | 780 | 82 | (2.07) |
| 1 | 1010±17 | 106±2 | (2.69±0.05) |
| 2 | 580±17 | 121±4 | (3.09±0.09) |
| 2 | 780 | 163 | (4.15) |
| 2 | 1010±17 | 212±4 | (5.37±0.09) |

Mesh was analyzed using a Phenom Pro Desktop scanning electron microscope (SEM, NanoScience) to capture fiber morphology, surface topography, and diameter. All samples were sputter coated with 5nm of gold before imaging. Fiber morphology was assessed for overall homogeneity and irregularity in morphologies from wet collection or multiple jets and captured using a 10 kV accelerating voltage, backscatter detector, and a magnification of 1000x. For fiber diameter and fiber fraction quantification, raster imaging was performed to obtain 5 images per mesh and each image was analyzed with ImageJ using the plugin DiameterJ. From the initial segmentation binary-colored segmented images produced with the algorithms M3, M5, M7, S2, S3, and S7 were used for determining fiber diameter and fiber diameter distribution. In the case of the plugin being unable to analyze one of the chosen segmentations another was chosen from the original set produced and substituted into the folder for analysis. From the same images, fiber fraction was calculated based on algorithm value output. All quantitative output for each tested condition has been collated for reference in **Table S1**.

### Statistical Analysis

For each group, five images from 8mm diameter specimens of mesh for 3 specimens per sample and 3 samples at each set of parameters were analyzed (n=9 specimens, 45 data points).

For plotting purposes, the 5 points representing individual punches were averaged to output a total of 15 points per group. Post-collection sample quantitative data was re-analyzed with a new ROUT outlier analysis (Q=1%) and single point outliers were pulled from each subset of data. Additionally, individual punches showing abnormal sample morphology in ≥3 SEM images were also removed from data sets (done post outlier analysis). Abnormal morphology was defined as visually webbing and splitting, clearly merged fibers, or general agglomeration of fibers visible throughout the image and due to abnormal jet morphology during the sample collection. In all data sets for groups with individual punches removed there is a marked sample size of n=8 and for un-marked groups a sample size of n=9.

All statistical tests and graphing were performed using Prism GraphPad software. Prior to any statistical analysis, data sets were tested for normality using a Shapiro-Wilk normality test. For all groups which passed the normality test, a Brown-Forsythe one-way ANOVA was used and for all groups which did not pass the normality test a non-parametric Kruskal-Wallis test was used. For normally distributed groups analyzed by Brown-Forsythe one-way ANOVA significance p is denoted using #. For groups analyzed by a non-parametric Kruskal-Wallis test significance p is denoted using *.

## Results

### Fiber Alignment

All groups were assessed for alignment and fiber properties via image-based analysis of SEM micrographs. **Figure 2** shows representative images of samples for every condition tested. In the representative images, there is general evidence of increasing alignment with increasing rotational speed for both the M3/4 and M1 samples. This is shown in **Figure 2A and 2B**. In the case of samples generated using the M2 mandrel, rotational speed appears to have limited additional effects on sample alignment with fibers that appear well aligned at nearly every set of tested parameters as is evident in **Figure 2C**. For each mandrel diameter, there appears to be at least one group of parameters which have a stronger influence on fiber alignment than an increased speed. This is particularly notable in the 30cm collection distance, and 18kV applied voltage groups. Micrographs of control groups collected on a stationary grounded plate and at identical processing parameter conditions can be seen in **Figure S2.**

**Figure 2.**
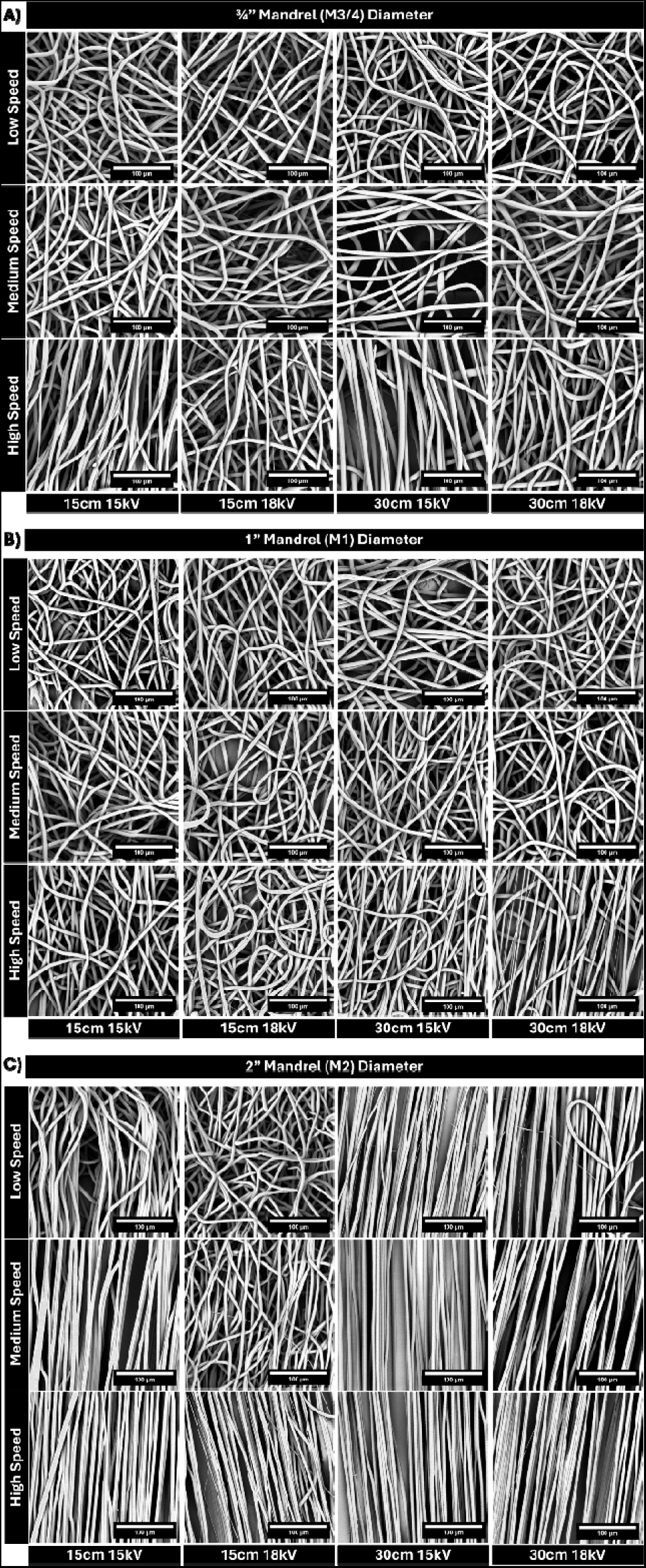
Representative SEM micrographs of samples at all conditions tested for A) ¾” diameter rotating mandrel, B) 1” diameter rotating mandrel, and C) 2” diameter rotating mandrel, all at 1000x magnification.

To offer a quantitative evaluation of fiber alignment, the full group of SEM micrographs for all parameters were analyzed using the ImageJ DiameterJ plug-in and this data is collated and visualized in **Figure 3**. In all groups tested, all speeds resulted in increased fiber alignment when compared to stationary controls; however, this increase was not always significant. Fiber alignment in all the sample groups at every tested parameter is highest in samples collected with the M2 mandrel. This is also evident in a revisualized representation of this data in **Figure S3**. The effect of increased alignment with increased mandrel diameter is only lessened in groups collected at lower speeds or at the lower collection distance with higher voltage or higher collection distance with lower voltage. In the 15cm, 18kV samples, there was relatively low alignment for each mandrel diameter at any given speed including the M2 mandrel groups which only showed high alignment at a high collection speed.

**Figure 3.**
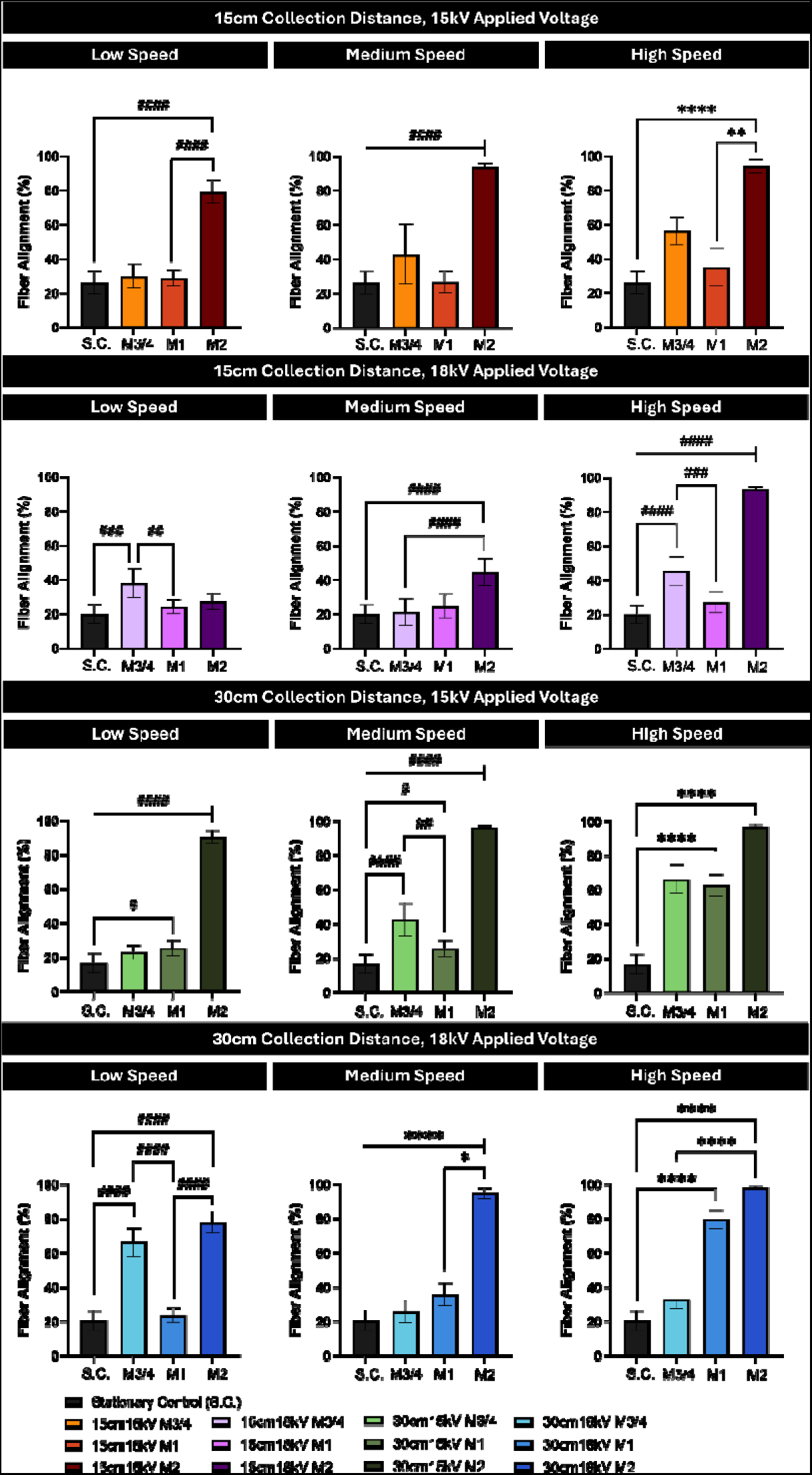
Average fiber alignment for all samples. (n=8 for M3/4 15cm15kV high speed, M3/4 15cm18kV medium speed, and M1 15cm15kV high speed, and n=9 for all other groups) #p≤0.0332, ##p≤0.0021, ###p≤0.0002, ####p≤0.0001. *p≤0.0332, **p≤0.0021, ***p≤0.0002, ****p≤0.0001

As shown in the ¾” diameter mandrel, an increased rotational speed typically increased alignment while in the M1 mandrel the collection voltage and distance appear to have much greater impacts on level of alignment. Similarly, despite nearly all conditions producing highly aligned samples on the M2 mandrel, processing at 18 kV applied voltage and 15 cm collection distance resulted in levels of alignment comparable to the unaligned controls. Overall, many of both the M1 and M3/4 diameter mandrel samples showed comparable levels of alignment to the unaligned stationary collection control samples (**Figure S4)**. In general, minimal changes in sample alignment were observed for samples collected at different speeds on a single diameter mandrel, especially with increased mandrel diameter. This trend can be seen in **Figure S4**.

### Fiber Fraction and Fiber Diameter

As shown in **Figure 4**, fiber fraction remained relatively consistent in all groups collected at the low and medium speed regardless of collector diameter. At a high collection distance, the samples generated on the M2 mandrel show significantly increased fiber fraction compared to both the stationary controls and smaller diameter mandrels. This trend is extended in the high-speed collection groups where differences in fiber fraction appear more distinct specifically in the 30cm collection distance group which shows increased fiber fraction with increased mandrel diameter.

**Figure 4.**
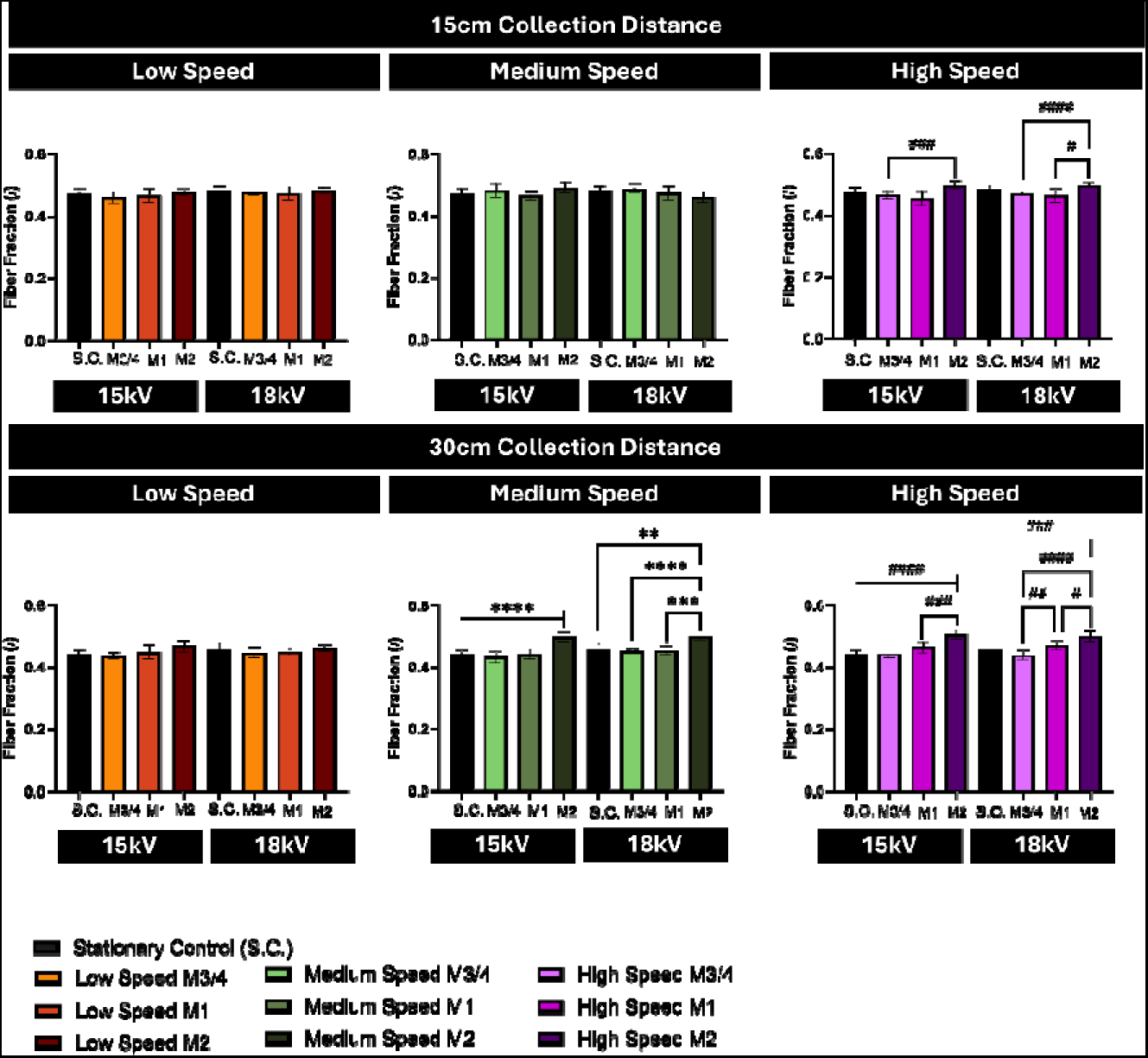
Average fiber fraction for all samples. (n=8 for M3/4 15cm15kV high speed, M3/4 15cm18kV medium speed, and M1 15cm15kV high speed, and n=9 for all other groups) #p≤0.0332, ##p≤0.0021, ###p≤0.0002, ####p≤0.0001. *p≤0.0332, **p≤0.0021, ***p≤0.0002, ****p≤0.0001

As shown in **Figure 5**, despite significant differences between many groups made using the M3/4 mandrel, fiber diameter of all groups ranged only from mean values of 5.1 μm-6.4 μm, a total average range of ~1.3 microns. In samples generated using the M1 mandrel, fiber diameter decreased with increased speed for nearly all conditions tested within a narrow range overall from mean values of 4.7 μm-6.9 μm, a total average range of ~2 microns. In samples generated using the M2 mandrel, fiber diameter decreased with increasing speed for nearly all conditions tested and ranged from a mean fiber diameter of 4.2 μm-5.7 μm, a total average range of ~1.5μm. In general, the mean fiber diameter of samples decreased with increasing mandrel diameter regardless of other processing parameters. In nearly all conditions, collection on a rotating mandrel decreased fiber diameter compared to stationary controls and this effect was more pronounced at higher collection distances and increased mandrel diameters.

**Figure 5.**
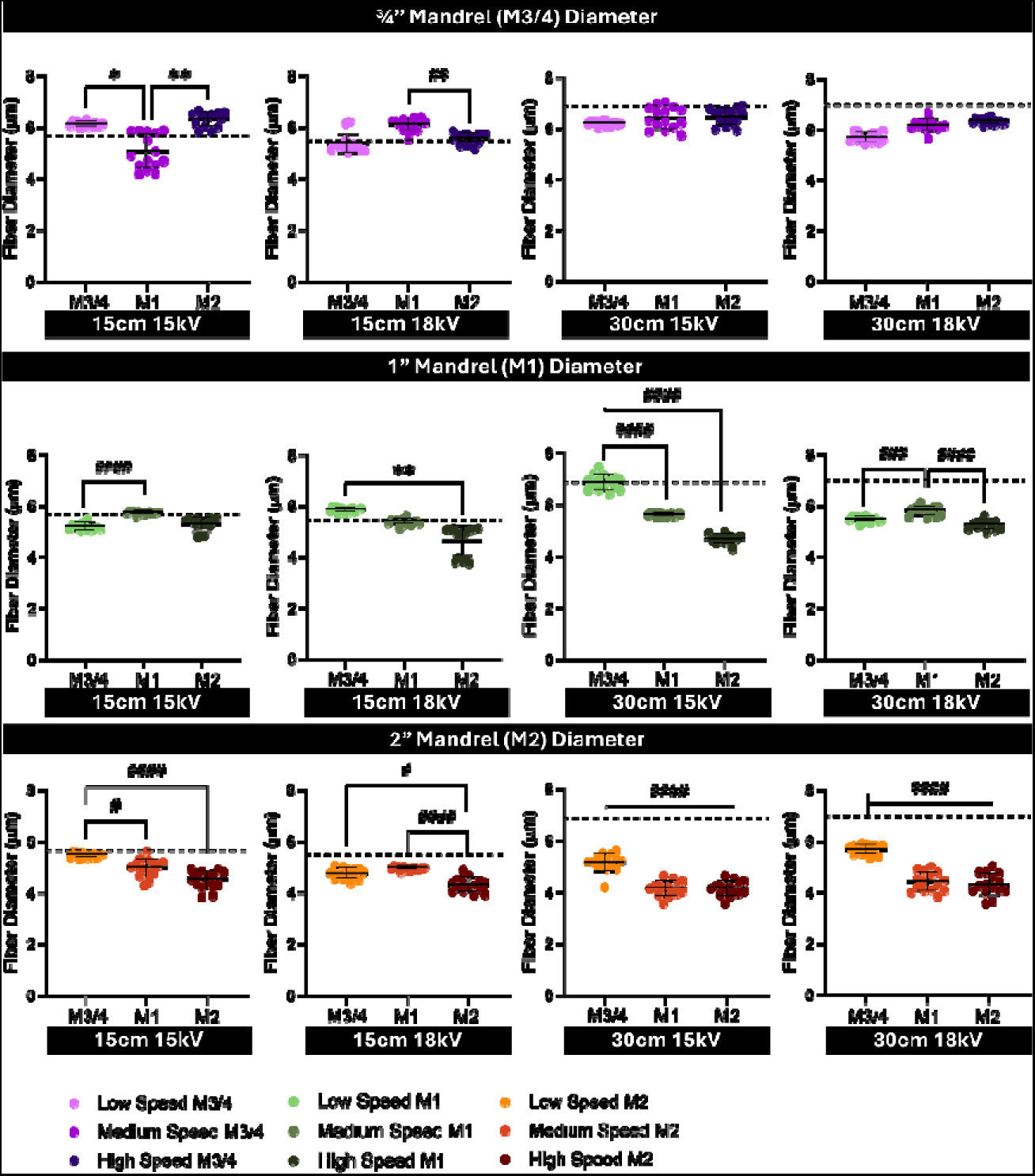
Average fiber diameter for all samples with dotted line representing stationary control values. (n=8 for M3/4 15cm15kV high speed, M3/4 15cm18kV medium speed, and M1 15cm15kV high speed, n=9 for all other groups) #p≤0.0332, ##p≤0.0021, ###p≤0.0002, ####p≤0.0001. *p≤0.0332, **p≤0.0021, ***p≤0.0002, ****p≤0.0001.

### Main Effects Summary of Importance

To demonstrate the importance of each processing parameter modulated in this study, a linear regression model was applied to each measured output (fiber diameter, fiber fraction, and fiber alignment) using the JMP software. As shown in **Figure 6**, the variable with the largest effect on all outcomes is the mandrel diameter followed by collection distance for fiber diameter and fiber fraction and rotational speed for fiber alignment. For all outcomes the applied voltage was assessed as having the least impact. Mixed effects are not included in this data analysis as the secondary mixed effects obscure contributions of the main study variables.

**Figure 6.**
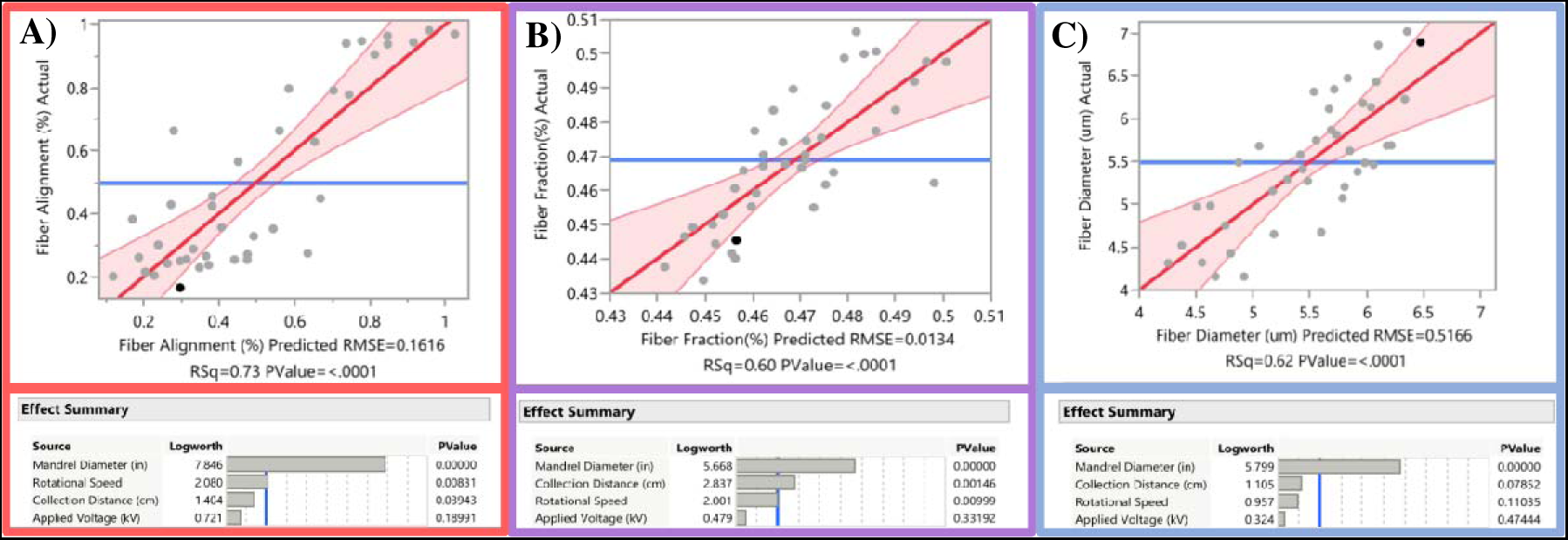
Main effects summary of importance and linear regression effects plot for mean fiber alignment **(A)**. mean fiber fraction **(B)**, and mean fiber diameter **(C)** of all analyzed sample groups.

## Discussion

Classically, increasing the rotational speed in a rotating mandrel electrospinning set-up increases collected fiber alignment. However, inconsistencies have been noted in this trend throughout the literature. This study demonstrates that the diameter of the rotating collector has a significant impact on fiber alignment at all speeds tested counter to what has been previously shown in other systematic studies.^11,27,28^ This difference in study outcomes can likely be attributed to the differing polymer systems, the scale of fiber diameters (nm vs μm), and the variance in parameters tested outside of mandrel diameter and rotational speed. Specifically, each other systems noted as examining these effects used thermoresponsive poly(N-isopropyl acrylamide) (PNIPAM)^27^, poly(ethylene terephthalate) (PET)^33^, and poly(−lactide-co-ε-caprolactone) (PLA-PCL)^35^. Other authors have noted that a better predictor of fiber alignment in samples is the linear velocity rather than rotational speed^36^ and the results of this study further emphasize this point. However, this study also emphasizes that while linear velocity is important, it is most beneficial when the overall system exhibits minimal jet whipping to interfere with the efficacy of the linear velocity. Further, the combination of an increased mandrel diameter and specific processing parameters can achieve comparable or increased levels of alignment as compared to solely increased rotational speed.

This study also demonstrates that it is possible to decrease the collector speed for aligned scaffolds by tuning parameters like collection distance and applied voltage. In the majority of systems, higher speeds range from 1000 RPM-8000 RPM^22,28–31,37–41^ with other noted speeds ranging from 10,000-33,000 RPM^27^. In this study, the maximum speed was 1010 RPM, illustrating the ability to achieve highly aligned fibers with orders of magnitude slower speeds which is much safer for the operator. Similarly to stationary electrospinning systems, there is an optimal set of parameters which can be tuned for different systems with specifically sized mandrels and specific outcome needs. Interestingly, even when samples collected using a mandrel showed comparable fiber alignment to stationary controls, the fiber diameter of a majority of samples was significantly different compared to their stationary controls. This can easily be attributed to the added system turbulence from any rotation of a collector, the applied mechanical force, and different collector geometry. However, it is of note for any study intending to create a randomly aligned control group with comparable fiber diameters.

A visual phenomenon noted in this study is the lack of a significant bending instability in many collected samples. This lack of a instability in highly aligned samples mimics the direct to collector type of jet found in melt electrowriting systems and corroborates with other studies illustrating that the ideal method of inducing alignment in samples is to create a collection type that removes or decreases the whipping region of jets.^11^ Based on the outcomes of highest alignment occurring for samples collected with the M2 mandrel, it is proposed that the dominant mechanism behind the alignment is a relatively weak electrical field which in combination with increased drawing power from the mandrel creates a stabilizing force for the electrospinning jet and a linear speed that matches or exceeds the extrusion speed of the solution. This might be mathematically explained through calculation of additional longitudinal strain rate applied by the added force of a rotating mandrel to calculations already established by Reneker et al.^42,43^ For any mandrel diameter below 2 inches (M2), results indicate that the dominating mechanism behind effective fiber alignment is an increased rotational speed of the collector. However, it is likely given these results that an optimized applied voltage and collection distance could allow for highly aligned fibers at decreased speeds.

For a set distance and rotational speed, the impact of voltage on fiber diameter is reduced by an increased mandrel diameter and this effect is enhanced by a larger collection distance. It has been shown that at higher collection distances, the overall decreased strength of electrical field increases time to collector^44^ which allows increased evaporation of solvent before collection.^45^ This increased evaporation results in an applied force on more solid jets which allows for less effective stretching of fibers that would likely cause a decrease in fiber diameter. However, for these samples, with increased collection distance on a larger mandrel diameter, fiber diameter is lower than those at lower collection distance as jets maintain some level of bending instability that allows a decrease in diameter.

At lower speed and decreased mandrel diameter, the electrical field appears to have greater effects on fiber diameter, and this can be attributed to a change in the dominating factors of system physics. Based on this study, it is likely that in these cases the electrical field dominates the system resulting in larger influence from the applied voltage. When comparing a set distance and voltage, increasing the speed typically decreases fiber diameter and this effect is enhanced by increased mandrel diameter. Not only do these results illustrate the importance of increased mandrel diameter, but they also further demonstrate that an increased mandrel diameter may create difficulty achieving a specific fiber diameter or porosity without further optimization for specific polymer systems. It is equally clear that in cases where a smaller diameter mandrel may be required, experimenters may rely more heavily on manipulation of solution and processing parameters like molecular weight and flow rate to achieve alignment at lower speeds.

Based on both trends in the literature and the polymer physics principles of increased packing and density with aligned chains, it was expected that there would be an increase in fiber fraction with alignment.^22^ Results showed some increase in fiber fraction between mandrel diameters indicating some increased level of packing and associated decrease in overall sample porosity with increased alignment of samples produced on larger mandrels. At the shorter collection times used for this study, the final sample thickness of highly aligned scaffolds was much lower than for unaligned scaffolds and this is evident both in scaffold appearance and in the SEM micrographs analyzed to obtain this final data. Even the minimal trend of increasing fiber fraction indicates an increased level of packing in aligned samples. Therefore, it is likely that samples collected at longer time scales would demonstrate a more distinct difference in sample fiber fraction, especially in samples with high levels of alignment.

Throughout this study, some interesting differences in fiber organization and therefore overall material alignment at different length scales was observed. It was noted that while many samples appeared visually aligned by eye, these same samples exhibited completely random fiber orientation upon analysis of scanning electron micrographs. While no analysis has been performed on the molecular chain orientation of these samples, the effects of both the electrospinning process and collection of aligned fibers through mechanical methods like rotating collectors on polymer molecular chain alignment have been shown.^13,32,33,36,37,46–51^ This is relevant to readers who may be concerned with specific mechanical properties of collected scaffolds and we suggest that there are three relevant scales of organization in these materials: macroscopic (visual), microscopic (SEM), and sub-micron (polymer chain). All these scales of varying organization may impact the final mesh properties and should be a relevant consideration in any experimental plan utilizing aligned electrospun materials.^52^

Limitations in interdependent variables in this study design are noted. For example, modulating rotational speed and mandrel diameter inherently alters linear speed. We chose to treat these as separate experimental parameters due to the other factors which may be modified by changes to a mandrel diameter including electrical field distribution and atmospheric turbulence at the mandrel surface. Lastly, compositional parameters including polymer molecular weight, solution viscosity, or solution conductivity were not modulated in this study. Future work will focus on characterizing the mechanical effects of increased alignment of fibers through differential scanning calorimetry, uniaxial mechanical testing, and x-ray diffraction to further elucidate changing crystallinity (i.e. polymer chain alignment) at a variety of length scales.

## Conclusions

This study successfully demonstrates the key parameters needed to induce a high level of alignment using a rotating mandrel system and impacts on fiber alignment and fiber fraction. In a single mandrel system, increased speed generally increases fiber alignment. However, this study has shown that an increased mandrel diameter is significantly more effective at increasing fiber alignment for low micron-scale electrospun fibers compared to speed for all tested groups. Further, the speeds used in this study are relatively low compared to commonly speeds suggesting that it is possible to obtain well-aligned fibers at these lower and safer operational speeds when modifying mandrel diameter or applied electrical field through manipulation of collection distance and voltage. This is important to note as other studies have shown that high collection speeds do not always increase alignment and can lead to breakage and inconsistent fiber diameter in scaffolds.^53^ Results from this study can be translated for other polymer-solvent systems and are applicable for any studies attempting to produce highly aligned electrospun fibers. Future work will focus on applying these system parameters to production of aligned nanofibers to elucidate whether the same governing principles dominate at a different length scale.

## Supporting Information

Data included in the supporting information includes an experimental parameter overview graphic (**Figure S1**), a collated table of all quantitative parameter outputs for each analyzed condition (**Table S1**), representative SEM micrographs of stationary control samples (**Figure S2**), and two plots of fiber alignment output with differing visual organizations (**Figure S3-4**).

