## Supplemental Figures 1-4 and Supplemental Table 1 for "Mandrel Diameter is a Dominating Parameter for Fiber Alignment Control in Rotating Mandrel Electrospinning Systems"

Address for all authors: 850 Republican St, Seattle, WA, 98109

Graphical Abstract:

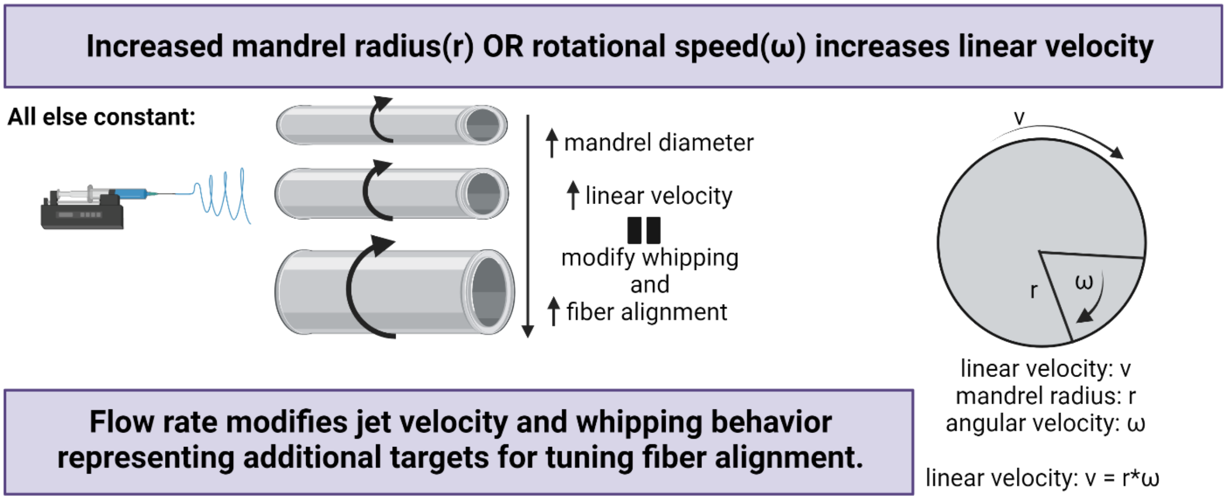

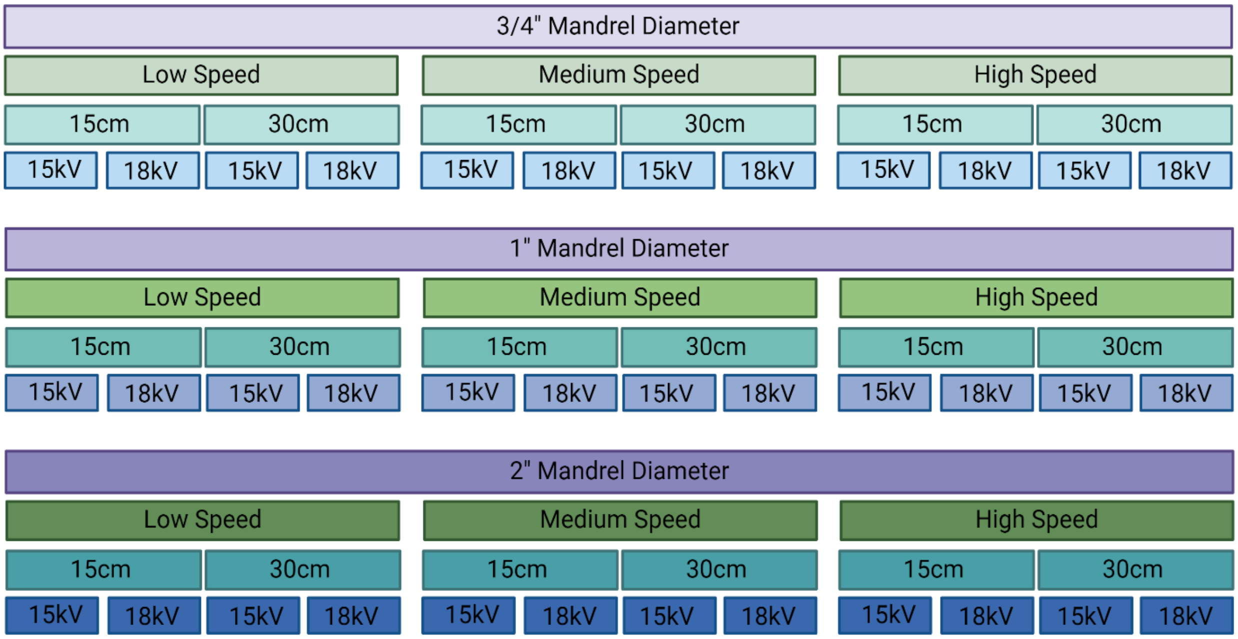


**Figure S1.** Experimental parameter overview separated by mandrel diameter

**Table S1.** Average fiber diameter, fiber fraction, and fiber alignment for all collected sample groups.

| Mandrel Diameter (in) | Collection Speed (RPM) | Collection Distance (cm) | Applied Voltage (kV) | Mean Fiber Diameter (µm) | Mean Fiber Fraction (/) | Mean Fiber Alignment (%) |
|---|---|---|---|---|---|---|
| 0 | 0 | 15 | 15 | 5.7±0.12 | 0.47±0.0 | 26±7 |
| 0 | 0 | 15 | 18 | 5.5±0.15 | 0.48±0.0 | 20±5 |
| 0 | 0 | 30 | 15 | 6.9±0.38 | 0.45±0.0 | 16±6 |
| 0 | 0 | 30 | 18 | 7.0±0.32 | 0.46±0.0 | 20±6 |
| 0.75 | 580±17 | 15 | 15 | 6.1±0.10 | 0.46±0.0 | 30±7 |
| 0.75 | 780 | 15 | 15 | 5.1±0.66 | 0.48±0.0 | 43±17 |
| 0.75 | 1010±17 | 15 | 15 | 6.3±0.27 | 0.47±0.0 | 56±8 |
| 0.75 | 580±17 | 15 | 18 | 5.4±0.35 | 0.48±0.0 | 38±8 |
| 0.75 | 780 | 15 | 18 | 6.1±0.24 | 0.49±0.0 | 21±8 |
| 0.75 | 1010±17 | 15 | 18 | 5.6±0.20 | 0.47±0.0 | 45±9 |
| 0.75 | 580±17 | 30 | 15 | 6.2±0.09 | 0.44±0.0 | 23±4 |
| 0.75 | 780 | 30 | 15 | 6.4±0.43 | 0.43±0.0 | 42±10 |
| 0.75 | 1010±17 | 30 | 15 | 6.5±0.32 | 0.44±0.0 | 66±8 |
| 0.75 | 580±17 | 30 | 18 | 5.7±0.19 | 0.45±0.0 | 66±8 |
| 0.75 | 780 | 30 | 18 | 6.2±0.22 | 0.45±0.0 | 25±7 |
| 0.75 | 1010±17 | 30 | 18 | 6.3±0.10 | 0.44±0.0 | 33±5 |
| 1 | 580±17 | 15 | 15 | 5.2±0.14 | 0.47±0.0 | 29±5 |
| 1 | 780 | 15 | 15 | 5.7±0.06 | 0.47±0.0 | 27±6 |
| 1 | 1010±17 | 15 | 15 | 5.3±0.26 | 0.46±0.0 | 35±11 |
| 1 | 580±17 | 15 | 18 | 5.9±0.08 | 0.47±0.0 | 24±4 |
| 1 | 780 | 15 | 18 | 5.4±0.11 | 0.48±0.0 | 25±7 |
| 1 | 1010±17 | 15 | 18 | 4.7±0.57 | 0.47±0.0 | 27±6 |

| | | | | | | |
|---|---|---|---|---|---|---|
| 1 | 580±17 | 30 | 15 | 6.9±0.29 | 0.45±0.0 | 25±5 |
| 1 | 780 | 30 | 15 | 5.6±0.05 | 0.44±0.0 | 25±5 |
| 1 | 1010±17 | 30 | 15 | 4.7±0.17 | 0.47±0.0 | 63±6 |
| 1 | 580±17 | 30 | 18 | 5.5±0.10 | 0.45±0.0 | 24±4 |
| 1 | 780 | 30 | 18 | 5.8±0.17 | 0.46±0.0 | 36±7 |
| 1 | 1010±17 | 30 | 18 | 5.3±0.17 | 0.47±0.0 | 80±5 |
| 2 | 580±17 | 15 | 15 | 5.5±0.09 | 0.48±0.0 | 79±7 |
| 2 | 780 | 15 | 15 | 5.0±0.34 | 0.49±0.0 | 94±2 |
| 2 | 1010±17 | 15 | 15 | 4.5±0.35 | 0.50±0.0 | 94±4 |
| 2 | 580±17 | 15 | 18 | 4.8±0.21 | 0.48±0.0 | 27±5 |
| 2 | 780 | 15 | 18 | 5.0±0.06 | 0.46±0.0 | 45±8 |
| 2 | 1010±17 | 15 | 18 | 4.3±0.27 | 0.50±0.0 | 94±1 |
| 2 | 580±17 | 30 | 15 | 5.2±0.34 | 0.47±0.0 | 90±4 |
| 2 | 780 | 30 | 15 | 4.2±0.28 | 0.50±0.0 | 96±1 |
| 2 | 1010±17 | 30 | 15 | 4.2±0.28 | 0.51±0.0 | 97±1 |
| 2 | 580±17 | 30 | 18 | 5.7±0.18 | 0.46±0.0 | 78±6 |
| 2 | 780 | 30 | 18 | 4.4±0.37 | 0.50±0.0 | 95±3 |
| 2 | 1010±17 | 30 | 18 | 4.3±0.43 | 0.50±0.0 | 98±1 |

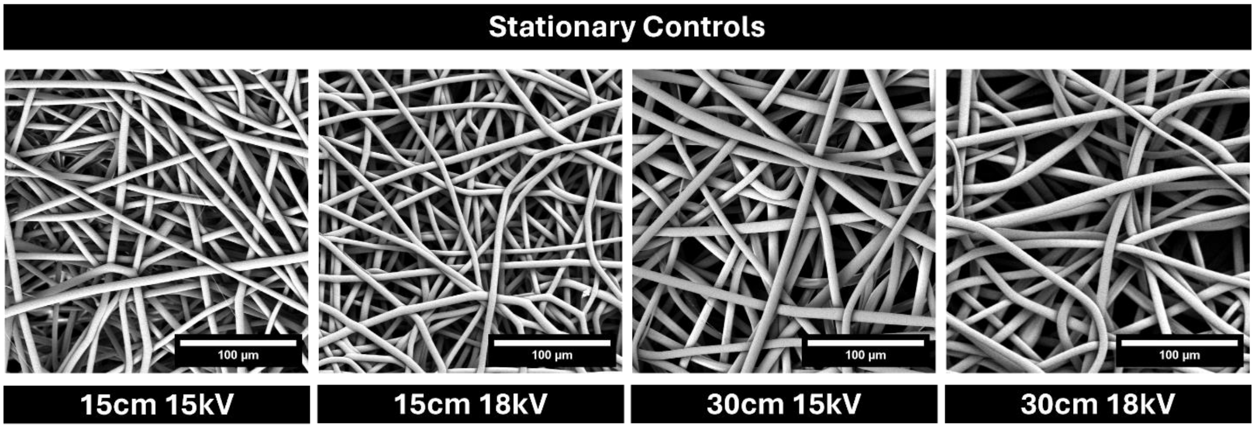


**Figure S2.** Representative SEM micrographs of all control samples collected on a flat, stationary copper plate.

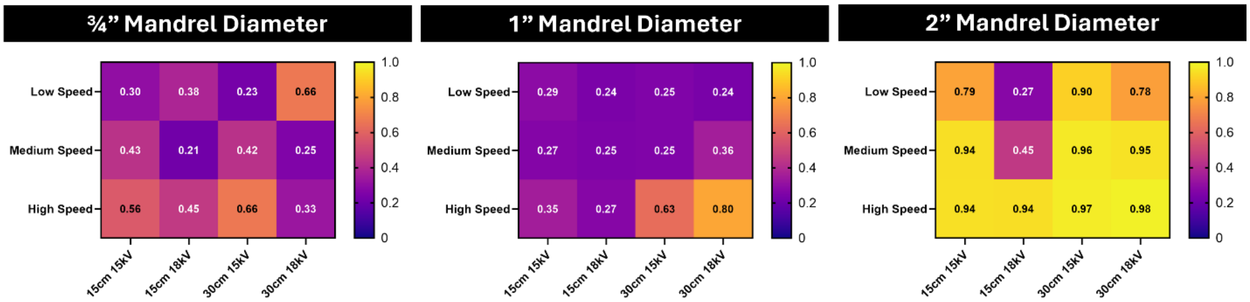


**Figure S3.** Heat maps of the average fiber alignment for all groups produced.

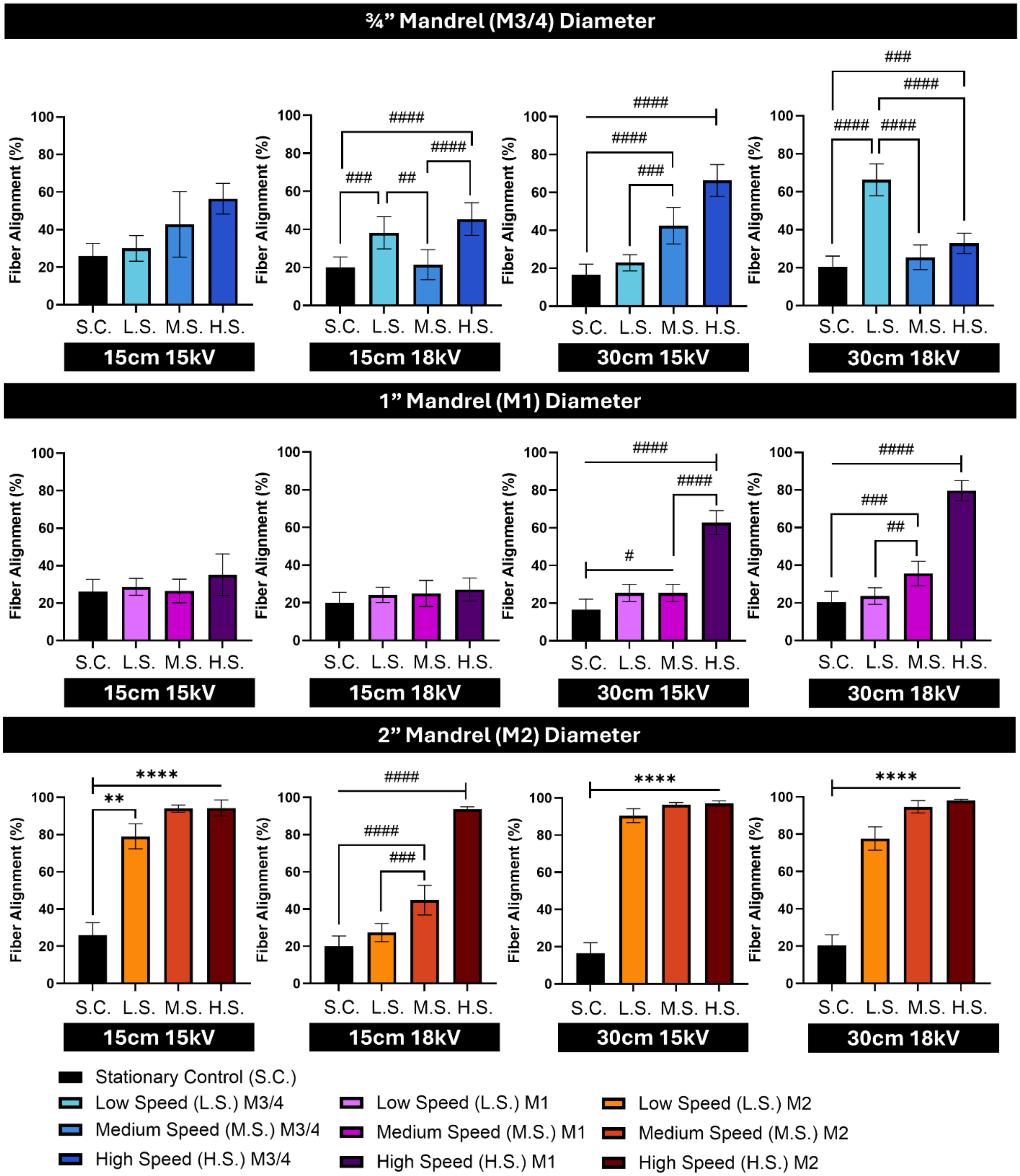


**Figure S4.** Average fiber alignment for all samples. (n=8 for M3/4 15cm15kV high speed, M3/4 15cm18kV medium speed, and M1 15cm15kV high speed, and n=9 for all other groups) #p≤0.0332, ##p≤0.0021, ###p≤0.0002, ####p≤0.0001. *p≤0.0332, **p≤0.0021, ***p≤0.0002, ****p≤0.0001
